# SUBCLINICAL ACUTE KIDNEY DISEASE: URINARY PROTEOMIC DIVERGENCE FOLLOWING NEPHROTOXIC AND ISCHAEMIC INJURY

**DOI:** 10.64898/2025.12.01.691515

**Authors:** Joana Mercado-Hernández, Isabel Fuentes-Calvo, David Martín-Calvo, Sandra M. Sancho-Martínez, Francisco J. López-Hernández, Carlos Martínez-Salgado

**Affiliations:** Translational Research on Renal and Cardiovascular Diseases (TRECARD), Department of Physiology and Pharmacology, University of Salamanca, 37007 Salamanca, Spain; Institute of Biomedical Research of Salamanca (IBSAL), 37007 Salamanca, Spain

**Keywords:** acute kidney injury, acute kidney disease, defective repair, urinary proteomic fingerprint, cisplatin, ischaemia

## Abstract

Defective renal repair following acute kidney injury (AKI) may lead to acute kidney disease (AKD), defined by persistent functional and/or structural abnormalities lasting up to three months. However, subclinical AKD often goes undetected due to the lack of specific diagnostic tools, as conventional biomarkers such as plasma creatinine (pCr) may remain within normal limits despite underlying structural damage.

To identify urinary proteomic signatures associated with subclinical AKD of differing etiologies, we induced AKI in Wistar rats using either toxic (cisplatin-induced) or ischemic (unilateral ischaemia-reperfusion, I/R) models, and monitored renal function, renal histopathology and urine composition over 42 days.

Renal function, as assessed by pCr and creatinine clearance, was rapidly and severely impaired following treatment with cisplatin and I/R, and normalized afterwards within 10 and 2 days, respectively. Despite functional recovery, histological analysis revealed persistent tissue injury—characterized by tubular dilation, necrosis, inflammation, and interstitial fibrosis—particularly in cisplatin-treated animals at day 42.

Urinary proteomic analysis identified 53 and 23 proteins that were differentially excreted with the urine in the cisplatin and I/R groups, respectively, compared to controls, and 11 proteins differed from one AKI etiology to the other. Additionally, 29 proteins appeared exclusively post-AKI, irrespective of cause. Most of these proteins are predominantly involved in the immune response, complement activation, haemostasis, and gluconeogenesis.

These findings suggest that urinary proteomic fingerprints may serve as sensitive indicators of subclinical AKD, reflecting underlying structural damage even in the absence of overt functional impairment. Such profiles could offer etiology-specific insights, while also enabling the early detection of AKD across diverse injury contexts.

## INTRODUCTION

Acute kidney injury (AKI) is characterized by a rapid decline in renal excretory function, clinically identified by an elevation in plasma creatinine (pCr) levels or the onset of oliguria [1]. AKI is associated with substantial morbidity and mortality, particularly within intensive care settings [2], and its incidence continues to rise, driven by an aging population and the increasing prevalence of comorbidities such as diabetes mellitus and hypertension [3]. Globally, AKI affects over 13 million individuals each year [4], contributing to nearly 2 million deaths annually [5]. In clinical practice, AKI is observed in approximately 21% of hospitalized patients, with incidence rates reaching up to 39% in critically ill patients admitted to intensive care units [6]. Although AKI is considered a transient syndrome, spontaneous recovery occurs in about two-thirds of the cases within 3 to 7 days, following the initial insult [7].

The kidney possesses a notable regenerative capacity, and in most cases, apparently full recovery following an episode of AKI can be achieved [8]. However, AKI is no longer regarded as a fully reversible condition. Even in mild cases where plasma creatinine (pCr) levels return to baseline, the risk of subsequent AKI episodes and mortality remains elevated [9]. This is partly attributed to maladaptive or aberrant repair processes, which prevent some regions of renal tissue from restoring their original architecture [9]. Incomplete recovery is associated with poorer clinical outcomes; however, recent evidence also suggests that even cases classified as fully recovered are linked to increased long-term mortality [10], irrespective of the severity of the initial insult—including moderate or subclinical AKI episodes [11] [12] [13] [14].

An additional consequence of incomplete or defective renal recovery is an increased risk of developing chronic kidney disease (CKD), particularly in the presence of comorbid conditions such as hypertension and diabetes mellitus [15]. However, prior to the clinical onset of CKD—or even in its absence—significant alterations in renal function and structure may persist for up to three months, without conforming to the traditional diagnostic criteria for either AKI or CKD. This intermediate state has been defined as acute kidney disease (AKD) [16]. Incomplete or defective renal repair represents a clinically relevant risk factor that remains difficult to diagnose, largely due to the inherent limitations of plasma creatinine (pCr), the universally accepted biomarker of renal function [3]. A measurable increase in pCr typically reflects only substantial reductions in glomerular filtration rate (GFR), which occur when a large number of nephrons are compromised. Moreover, due to compensatory mechanisms such as the recruitment of renal functional reserve, early or mild nephron injury may not result in detectable changes in GFR beyond the normality range [17]. During the recovery phase of AKI, pCr values may normalize even in the presence of persistent structural damage or incomplete restoration of renal function [18]. Previous work from our research group has demonstrated the presence of subclinical sequelae following apparent recovery from cisplatin-induced AKI, based on normalized pCr levels. These hidden alterations correlate with the development of undetected, subclinical AKD and are associated with an increased susceptibility to subsequent AKI episodes [19] [20].

Given that plasma creatinine (pCr) is not a reliable biomarker for detecting defective renal repair—and thus, for identifying subclinical or hidden AKD—there is a pressing need for novel diagnostic tools capable of detecting and monitoring these pathological changes. Among the most promising approaches is the identification of alternative biomarkers that reflect underlying pathophysiological processes. Our research group has previously identified several urinary biomarkers—including kidney injury molecule-1 (KIM-1), albumin, transferrin, insulin-like growth factor-binding protein 7 (IGFBP7), and tissue inhibitor of metalloproteinases-2 (TIMP-2)—which can detect subclinical sequelae following cisplatin-induced AKI [19] [20]. However, these biomarkers appear to be specific to nephrotoxic injury and may not be suitable for identifying structural damage associated with AKI of different etiologies. Moreover, many of these markers are also implicated in other renal pathologies, such as susceptibility to AKI [21] and progression to CKD [22], thereby limiting their diagnostic specificity. Therefore, it is essential to identify novel urinary proteomic fingerprints with differential diagnostic potential for detecting defective repair associated with AKD, particularly those induced by AKI of diverse etiological origins.

In the present study, we identified urinary biomarker fingerprints associated with subclinical AKD following the apparent recovery of renal function, using differential proteomic analysis in two experimental models of AKI in Wistar rats: a nephrotoxic model induced by cisplatin administration and a unilateral ischemia-reperfusion (I/R) model.

## METHODS

All reagents were purchased from Merck (Madrid, Spain), except where otherwise indicated.

### In vivo experimental model

Animals were treated in accordance with the ARRIVE guidelines and the Principles of the Declaration of Helsinki and the European Guide for the Care and Use of Laboratory Animals (Directive 2010/63/UE) and Spanish national and regional regulations (Law 32/2007/Spain, RD 1201/2005 and RD 53/2013). The Bioethics Committee of the University of Salamanca and the Regional Government of Castile and Leon, Ministry of Agriculture and Livestock approved all procedures for Animal Care and Use (institutional animal protocol 143-USAL).

Male Wistar rats (200–250 g; Charles River, Madrid, Spain) were maintained under controlled environmental conditions, with access to water and standard chow *ad libitum*. Rats were divided into three experimental groups (n=12 per group): cisplatin group (cPt), rats receiving cisplatin (5 mg kg^−1^, i.p.) at the beginning of the study; ischaemia/reperfusion group (I/R), rats subjected to one hour of warm renal ischaemia on the left kidney; control group (C), rats receiving saline solution (0.9% NaCl, i.p) and undergoing a SHAM surgery on the same days than the previous groups (Figure 1). For the renal I/R, rats were anesthetized with a mix of 80 mg/kg ketamine and 20 mg/kg xylazine. Through a medial laparotomy, the left renal artery was gently exposed, cleaned from surrounding fat, and clamped for 60 minutes, after which the clamp was removed to allow for reperfusion. The incisions were sutured, and a protocol of analgesia with 30 μg/kg buprenorphine was followed for the next 5 days at 12-hour intervals.

**Figure 1.**
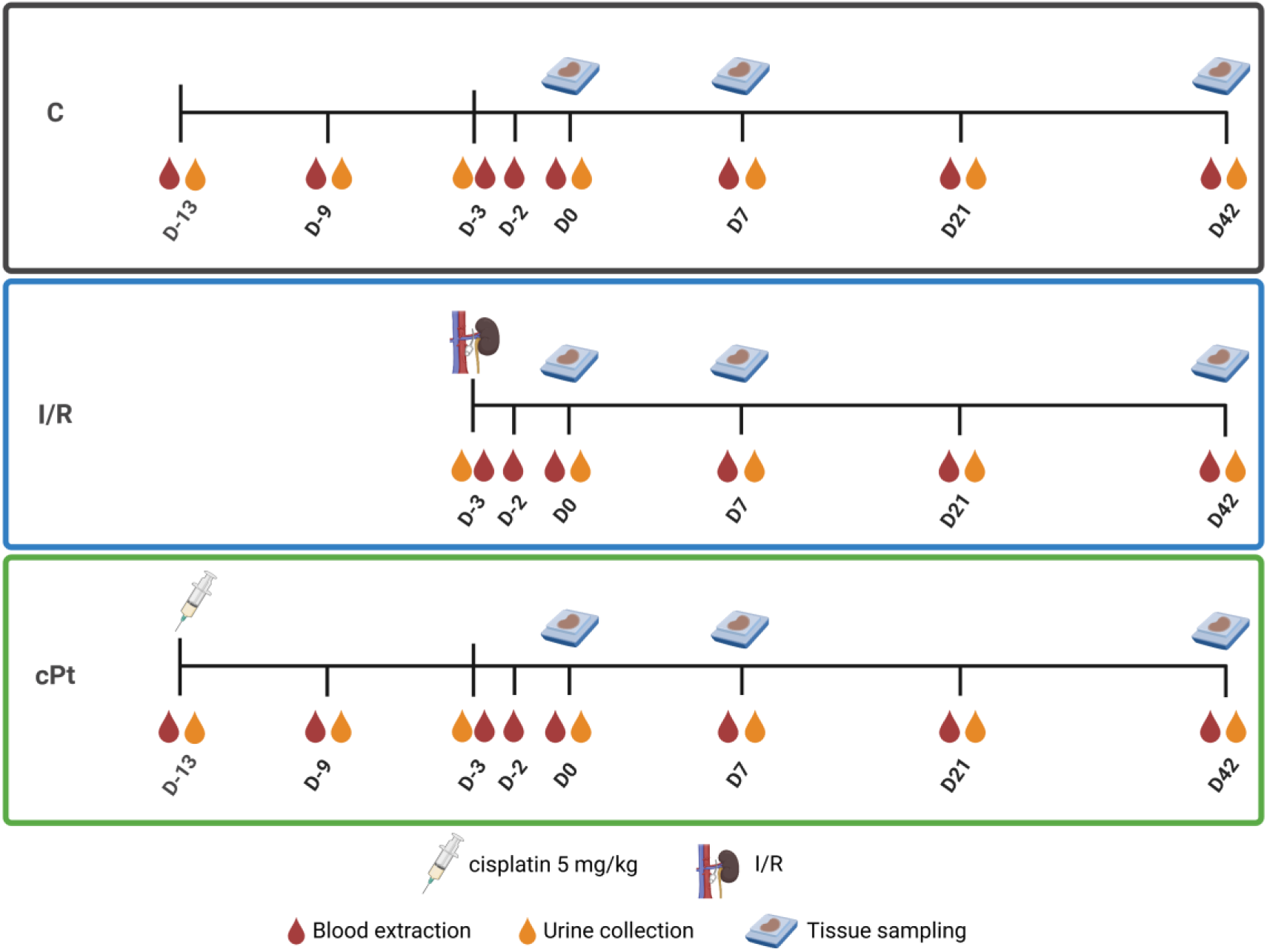
Time diagram of the in vivo experimental protocol and days for urine, plasma and tissue samples. D-13 (day -13), immediately prior to cisplatin administration; D-9, D-2 (days -9, -2), days of cisplatin or ischaemia-induced maximum kidney damage, respectively (highest pCr level); D-3 (day -3, day of ischaemic surgery) D0, day of recovery (pCr returns to basal levels); and D7, D21, D42 (days 7, 21, 42 post-recovery). I/R: ischaemia-reperfusion group.

### Sample collection

At selected time points (Figure 1), urine and plasma samples were collected to evaluate renal function. For urine collection, rats were individually allocated in metabolic cages. 24-h urine was collected, centrifuged, and stored at −80 °C. 600 μl blood was drawn from the tail vein, centrifuged and plasma was stored at −80 °C. Immediately before sacrifice at day 42 (50 mg kg^−1^ sodium pentobarbital i.p.), kidneys were perfused through the aorta with saline solution and dissected. One-half was frozen in liquid nitrogen and subsequently kept at − 80 °C. The other half was fixed in buffered 3.7% p-formaldehyde for histological studies.

Renal function was evaluated in the following days: D-13 (day -13), immediately prior to cisplatin administration; D-9, D-2 (days -9, -2), days of cisplatin or ischaemia-induced maximum kidney damage, respectively (highest pCr level); D-3 (day -3, day of ischaemic surgery) D0, day of recovery (pCr returns to basal levels); and D7, D21, D42 (days 7, 21, 42 post-recovery). Tissue samples were collected on D0, D7 and D42 for histological analysis.

### Histological studies

Upon surgical resection, kidney samples were fixed in 3.7% formaldehyde and embedded in paraffin blocks. 3 μm-thick slices were stained with haematoxylin-eosin and Masson’s trichrome. Microphotographs were obtained using DotSlide virtual microscopy technique under an Olympus BX51 microscope (Olympus, Tokyo, Japan). In a standardized manner, representative pictures of each sample were taken with an Olympus DP70 digital camera (Olympus, Tokyo, Japan). Five pictures were taken across the renal cortex and another five across the external medulla.

### Renal function assessment

Plasma (pCr) and urinary creatinine (uCr) concentrations were assessed using commercial kits based on the Jaffe reaction (QuantiChrom Creatinine Assay Kit, BioAssay Systems, Hayward, CA, USA). Glomerular filtration rate (GFR) was estimated by the creatinine clearance (ClCr), using the following formula: ClCr = uCr x UF / pCr, were UF is the urine flow. GFR was also estimated with the ACLARA equation, based on plasma creatinine and body weight values [23]. Plasma urea concentrations were determined using a commercial kit based on the improved Jung method (QuantiChrom Urea Assay Kit, BioAssay Systems). Urinary and plasma sodium were determined using a LAQUATWi B772 meter (Horiba, Kyoto, Japan). Proteinuria was quantified with a commercial kit based on the Bradford assay, following the manufacturer’s instructions (BioAssay System).

### Proteomic analysis

Five urine samples of each experimental group from day 42 were selected for proteomic analysis. An equivalent of 100 μg of total protein (corresponding to 60-400 μL of urine depending on the sample) was precipitated using cold methanol/chloroform, washed, and solubilized in 8M urea. After reduction and akylation, proteins were digested with 1:50 trypsin at 37 °C overnight. Peptides were acidified with TFA 3% and purified using reversed-phase chromatography with POROS 20 R2 resin. Peptide concentrations were determined using a QuBit 3.0 Flex Fluorometer (Thermo Fisher Scientific, Madrid, SpainFor LC-MS/MS analysis, 0.5 µg of peptides were loaded onto a PepMap100 C18 trap column (Thermo Fisher Scientific) and separated on an analytical PepMap RSLC C18 column (50 cm × 75 µm, 2 µm particle size, 100 Å pore size) using a Vanquish Neo IFC system (Thermo). Peptides were eluted with a 120-min linear gradient (2–35% ACN, 0.1% formic acid) at 250 nL/min.

Eluted peptides were ionized by positive electrospray ionization and analyzed on a Q Exactive HF mass spectrometer (Thermo Fisher Scientific) operated in data-dependent acquisition (DDA) mode. Full MS scans were acquired at a resolution of 60,000 (at m/z 200) across an m/z range of 350–1800, using an ionization voltage of 2.1 kV and a capillary temperature of 275 °C. The top 15 precursor ions (charge 2+ to 5+) were fragmented by higher-energy collisional dissociation (HCD) with a normalized collision energy (NCE) of 28%. MS/MS spectra were acquired at 15,000 resolution, and dynamic exclusion was applied for 30 s to avoid repeated precursor selection.

### Proteomics data processing

Mass spectral data were processed on Proteome Discoverer software v2.5 (Thermo Fisher Scientific). Spectra were searched with MASCOT against the Rattus norvegicus UniProt database (downloaded 10 January 2020; 29,850 entries). Search parameters included a precursor mass tolerance of 10 ppm, fragment tolerance of 0.02 Da, and up to two missed cleavages. Carbamidomethylation of cysteine was set as a fixed modification, while oxidation of methionine and protein N-terminal acetylation were considered variable modifications. Label-free quantification (LFQ) was performed based on precursor ion intensities normalized to total peptide abundance per sample. Only proteins identified with at least two unique peptides, |log₂ (abundance ratio)| ≥ 1, and a false discovery rate (FDR) < 0.05 were considered differentially expressed. Statistical significance was assessed by one-way ANOVA followed by Student’s t-tests, with p-values corrected by the Benjamini–Hochberg method.

The Differentially Expressed Proteins (DEP) list was used to generate a network of predicted associations using STRING database as previously described [24]. This network was further interrogated using the Reactome database to identify the biological process in which the proteins were involved.

### Statistical analysis

Renal function statistical analysis was performed using the GraphPad Prism 7 software (San Diego, CA USA). Data normal distribution was evaluated with the Shapiro–Wilk normality test. Data are shown as mean ± standard error of the mean. Two-way ANOVA with Šidák’s test were performed to compare experimental groups over time. A p value < 0.05 was considered statistically significant. Proteomics statistical analysis was performed with R (version 4.3.1; R Core Team, Boston MA, USA) using Analysis of Variance (ANOVA) to evaluate differences between conditions, followed by the Student’s t-test for significant comparisons (p value < 0.05). Non-parametric data were analysed using the 2 -tailed Mann-Whitney U test (function wilcox.test). In addition, the Benjamin-Hochberg correction was applied to control for FDR to obtain an adjusted p value.

## RESULTS

### Renal filtration apparently normalizes after AKI episodes of different etiology

Both cisplatin- and ischaemia-induced AKI result in significant elevations in plasma creatinine (pCr) and plasma urea nitrogen, accompanied by reductions in both measured and estimated creatinine clearance (ClCr) on days -9 and -2, respectively. However, by 34 days post-cisplatin administration and 23 days following ischaemia-induced AKI, pCr and ClCr values return to levels within the range considered non-pathological (Figure 2).

**Figure 2.**
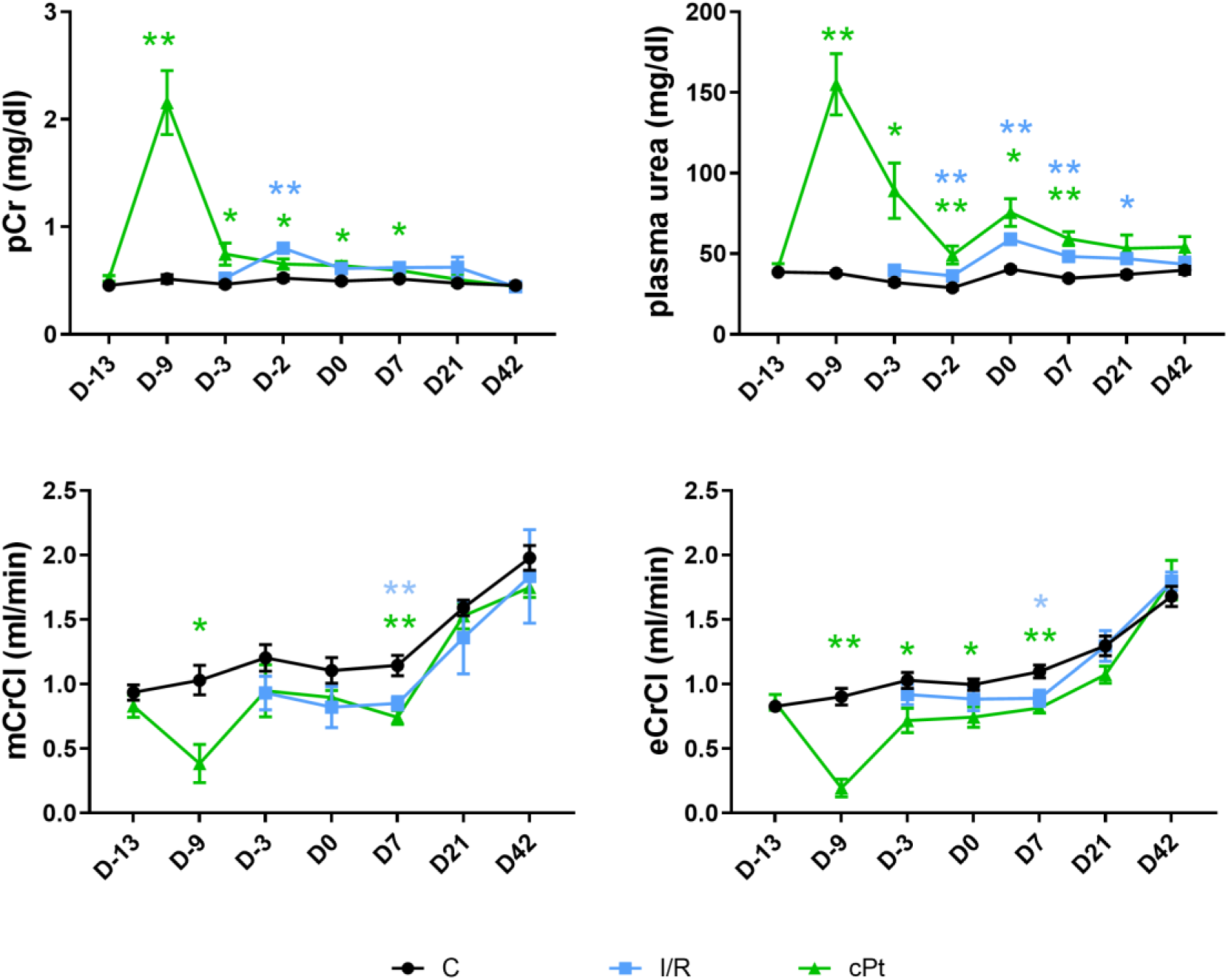
Renal function. Evolution of plasma creatinine concentrations, plasma urea, creatinine clearance (as an estimation of glomerular filtration rate) and creatinine clearance estimated by ACLARA equation, over the course of the experiment. Data represent the mean ± SEM of n = 9 per group. Two-way ANOVA with Šidák’s test were performed to compare the cisplatin-ischaemia group with the control group. * p < 0.05 and ** p < 0.01 versus control group in the same day. C: control; cPt: cisplatin group; eCrCl: estimated creatinine clearance (ACLARA formula); I/R: ischaemia-reperfusion group; mCrCl: measured creatinine clearance; D4: day of maximum kidney damage after cisplatin treatment; D-13, D-9, D-3, D0, D7, D21, D42: days -13, -9, -3, 0, 7, 21, 42 respectively.

### The kidney shows structural sequelae after functional recovery from AKI

AKI episodes induce structural alterations in the renal tissue that progressively ameliorate over time. In the I/R group, marked tubular dilation is observed on day 13 in both the renal cortex (Figures 3 and 4) and outer medulla (Supplementary Figures 1 and 2), accompanied by partial epithelial disorganization, the presence of cellular debris, and hyaline cast formation. By day 20, a reduction in the number of dilated tubules is evident, particularly in the cortex, along with early signs of tubular repair and reorganization (Figure 3, Supplementary Figure 1). However, this phase also coincides with an increase in extracellular matrix (ECM) deposition (Figure 4, Supplementary Figure 2). At one-month post-injury, most tubular abnormalities are resolved, although cortical fibrosis persists.

**Figure 3.**
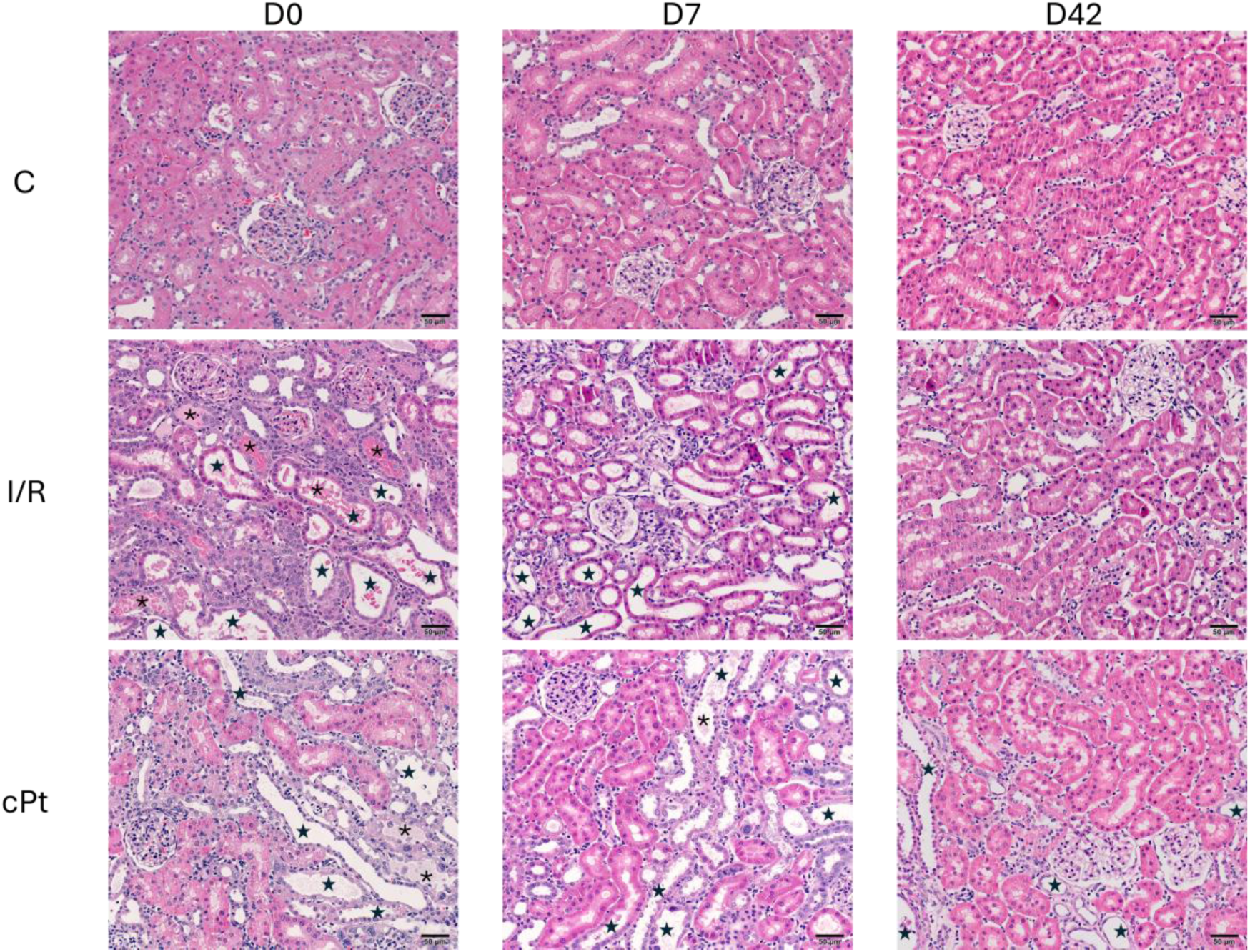
Renal histology. Representative images of renal cortex specimens stained with haematoxylin and eosin (HE) at days 0, 7 and 42 (magnification: 200X). Casts and cellular debris (*), tubular atrophy (star) and inflammatory infiltrates (#) are visible in both AKI groups, both in the cortex and in the outer medulla. This damage gradually decreases until day 42, where only isolated lesions are observed in the cPt group. C: control; cPt: cisplatin group; I/R: ischaemia-reperfusion group; D0, D7, D42: days 0, 7 and 42 respectively.

**Figure 4.**
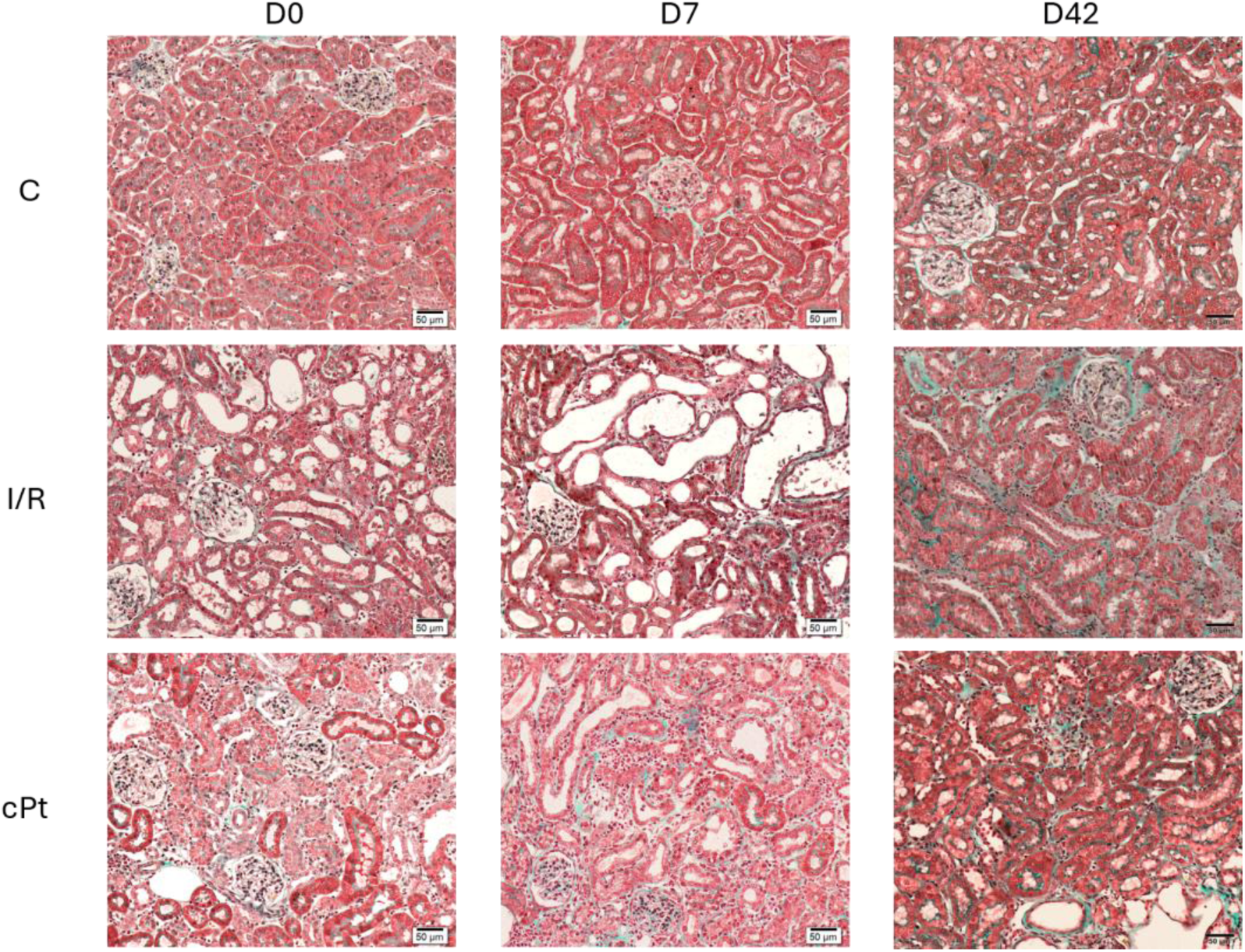
Fibrosis assessment. Representative images of renal cortex specimens stained with Masson trichrome at days 0, 7 and 42; magnification: 200X. There is a progressive fibrosis marked by the accumulation of collagen fibers (stained in green). C: control; cPt: cisplatin group; I/R: ischaemia-reperfusion group; D0, D7, D42: days 0, 7 and 42 respectively.

In the cisplatin-treated group, histological analysis on day 13 reveals pronounced epithelial disruption, inflammatory cell infiltration, increased ECM deposition, tubular dilation, and accumulation of intratubular cellular debris (Figure 3, Supplementary Figure 1). While partial structural recovery is observed at later time points—i.e., resolution of most acute tubular changes—fibrotic remodeling remains evident (Figure 4, Supplementary Figure 2). Notably, the extent of recovery in the cisplatin group is markedly less pronounced than that observed in the I/R group by the end of the experimental period.

These findings indicate that progressive fibrotic remodeling can occur in the context of subclinical AKD, under an apparently normal renal function.

### The proteomic fingerprint associated with subclinical AKD is different depending on the etiology of renal damage

Proteomic analysis at day 42 post-recovery reveals distinct urinary protein fingerprints depending on the etiology of the initial renal injury. In the cisplatin-treated group, the urinary excretion of 44 proteins showed significant differences compared to controls, with 8 proteins displaying decreased levels and 36 showing increased urinary excretion (Figure 5). Additionally, 9 proteins were exclusively detected in the urine of cisplatin-treated rats, absent in both control and ischemia groups (Table 1). These proteins are primarily associated with biological processes such as coagulation and haemostasis, immune response, and inflammation.

**Figure 5.**
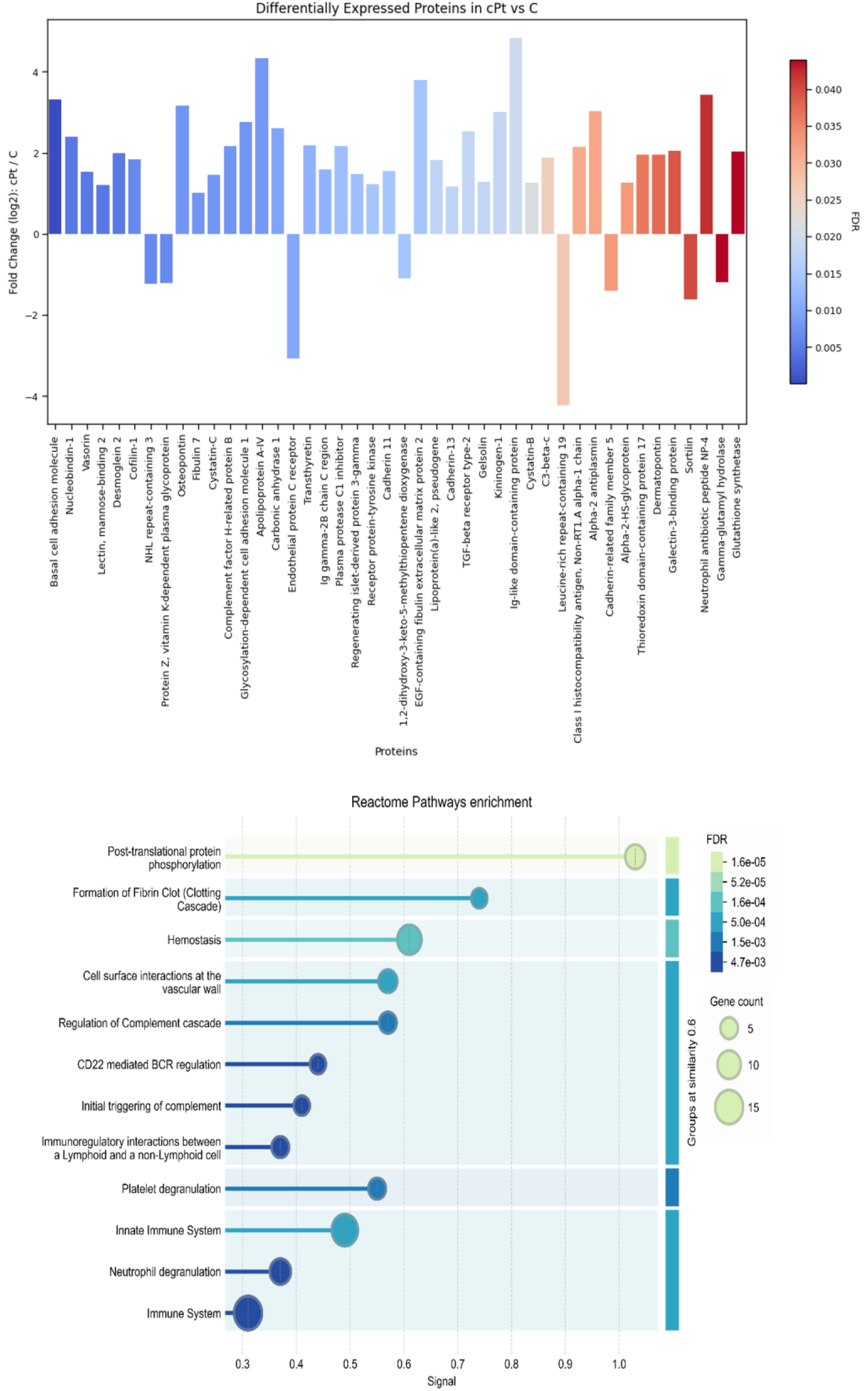
Differentially expressed urinary proteins in cisplatin vs. control group. Up-regulated and down-regulated urinary excretion of proteins between cisplatin and control groups at day 42 post-recovery, and biological processes associated with differentially excreted urinary proteins, as identified through Reactome Pathway enrichment analysis. Proteins unique to this comparison are highlighted in bold. C: control; cPt: cisplatin; FC: fold change; FDR: false discovery rate

In the ischemia group, 15 proteins exhibited differential urinary excretion compared to the control group, including 3 with reduced and 12 with elevated levels (Figure 6). Moreover, 8 proteins were uniquely detected in the I/R group but absent in both control and cisplatin-treated animals (Table 2). Functional enrichment analysis indicates their involvement in immune response and haemostatic processes.

**Figure 6.**
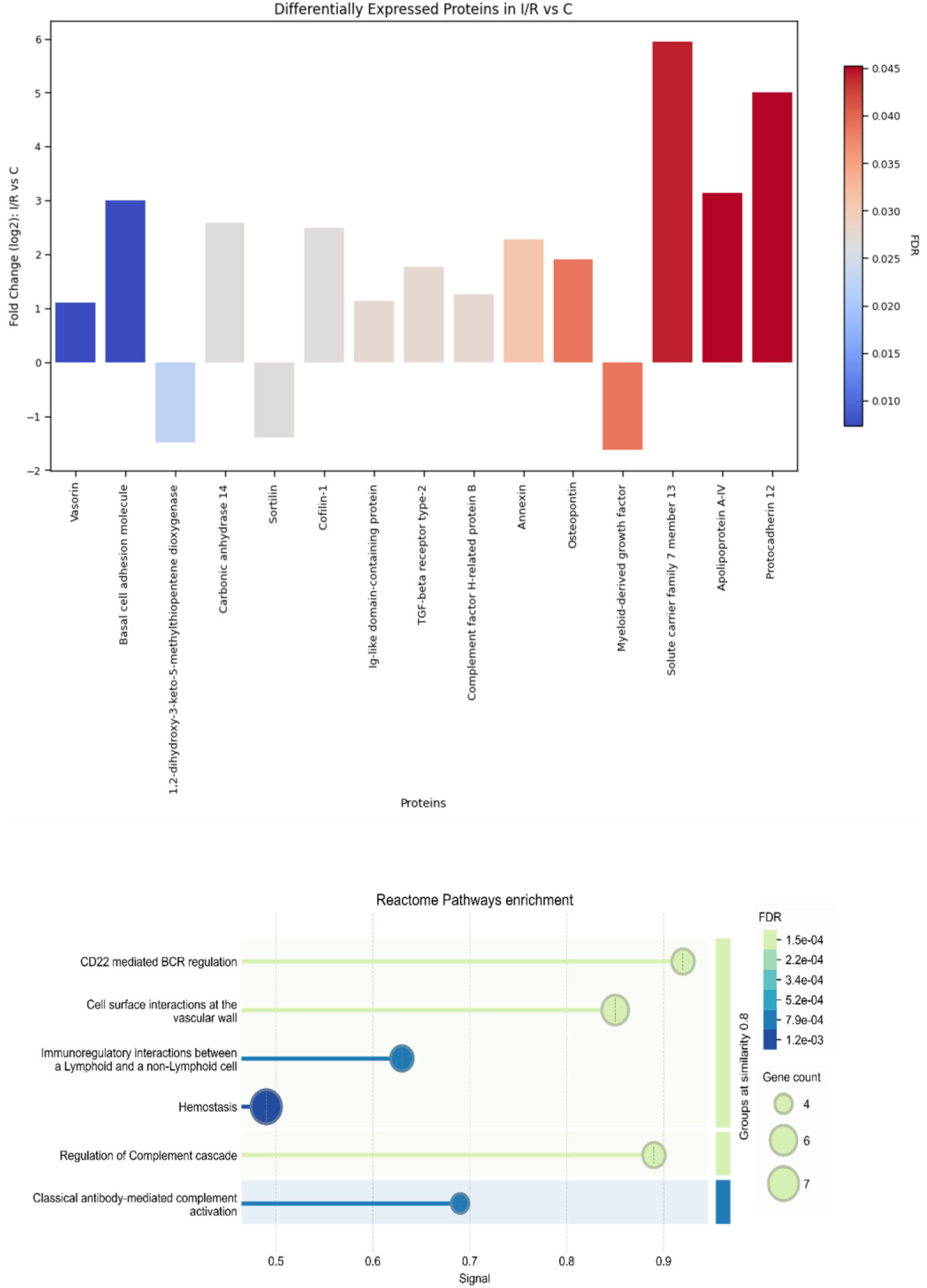
Differentially expressed urinary proteins in ischaemia vs. control group. Up-regulated and down-regulated urinary excretion of proteins between ischaemia and control groups at day 42 post-recovery and biological processes associated with differentially excreted urinary proteins, as identified through Reactome Pathway enrichment analysis. Proteins unique to this comparison are highlighted in bold. C: control; FC: fold change; FDR: false discovery rate; I/R: ischaemia group

Furthermore, 29 proteins were commonly detected in the urine of both the cisplatin and ischemia groups but were absent in controls. Of these, 17 showed increased and 12 showed decreased excretion in the cisplatin group compared to the ischemia group (Table 3, Figure 7). Additionally, 5 proteins present in control animals were undetectable in both injury models (Table 4). The proteins shared exclusively by the kidney injury groups are functionally linked to immune response and haemostatic regulation.

**Figure 7.**
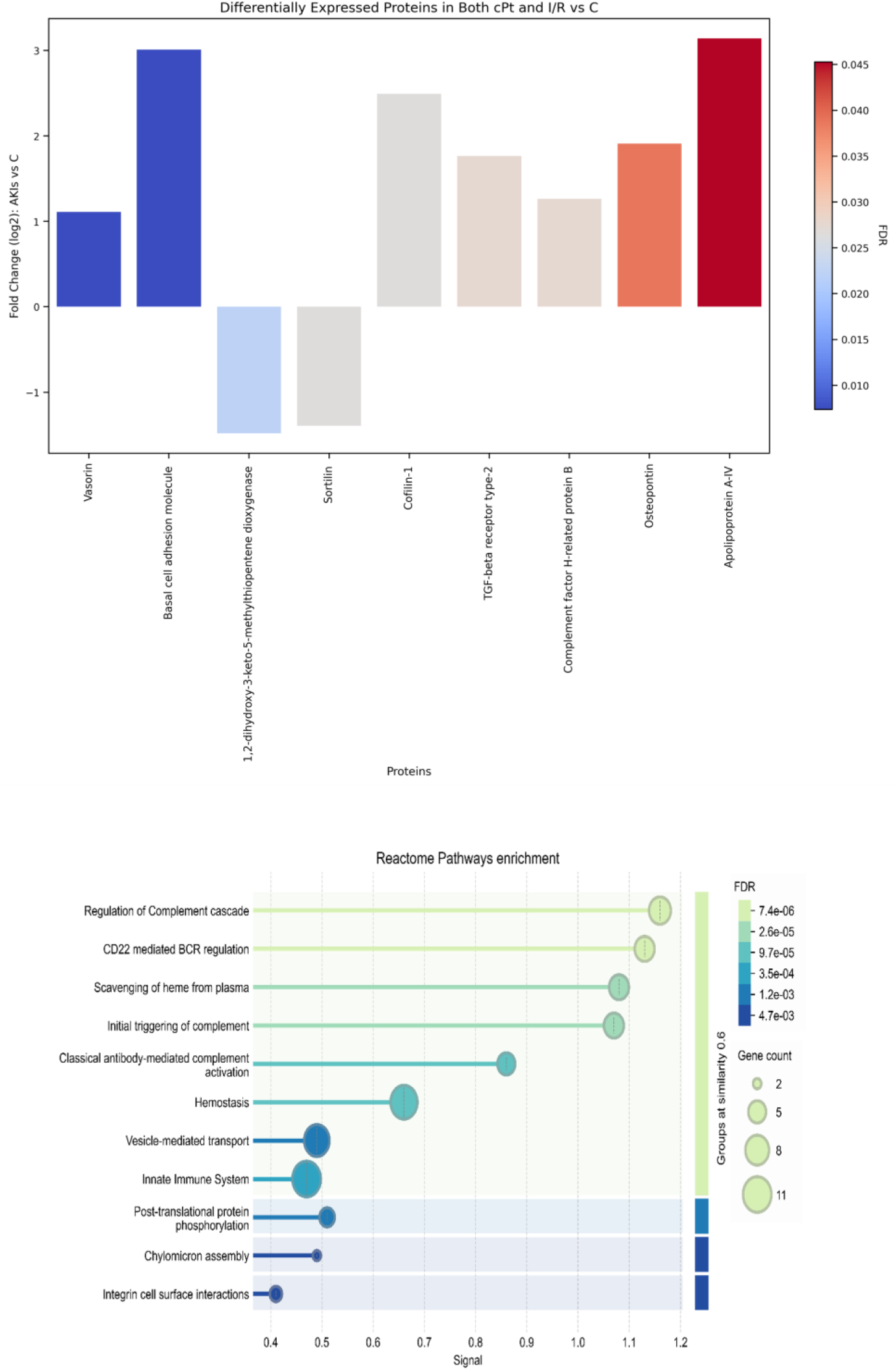
Differentially expressed urinary proteins in cisplatin + ischaemia (AKI) vs. control group. Up-regulated and down-regulated urinary excretion of proteins between cisplatin + ischaemia and control groups at day 42 post-recovery and biological processes associated with differentially excreted urinary proteins, as identified through Reactome Pathway enrichment analysis. Proteins unique to this comparison are highlighted in bold. C: control; cPt: cisplatin; FC: fold change; FDR: false discovery rate; I/R: ischaemia group

In summary, 44 proteins were differentially expressed between the cisplatin and control groups, with 33 being unique to this comparison. Of the 15 proteins differing between the ischemia and control groups, 6 were unique, while 9 of the 11 proteins distinguishing the cisplatin and I/R groups were exclusive to that comparison (Figure 8).

**Figure 8.**
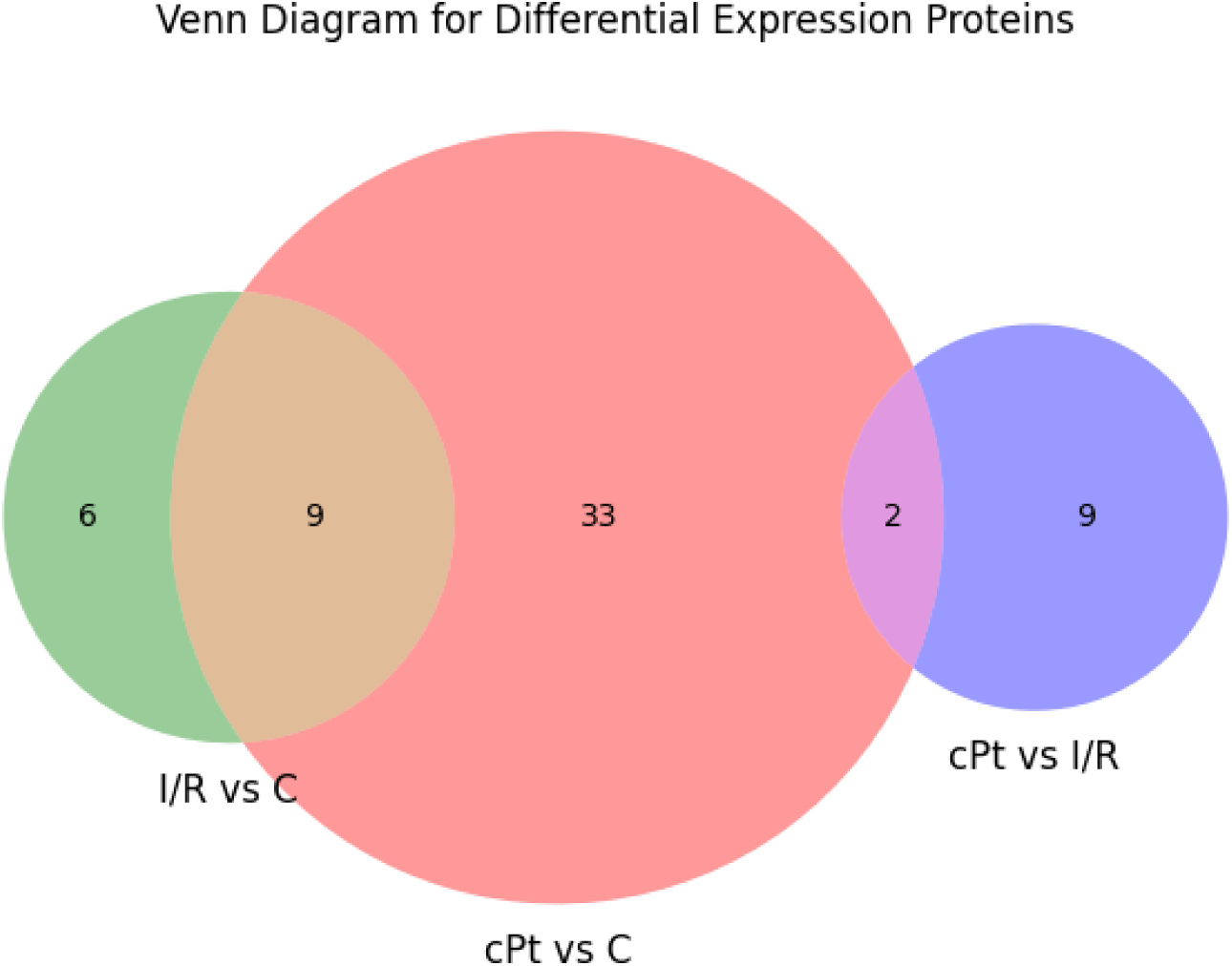
Venn diagram of differentially expressed proteins. The diagram shows the comparisons between the cisplatin, ischaemia, and control groups. cPt: cisplatin; I/R: ischaemia group

## DISCUSSION

The present study demonstrates that, although renal function appears to recover after episodes of AKI induced by either cisplatin administration or ischemia-reperfusion, structural and molecular alterations persist, indicating the presence of subclinical AKD. One of the strategies contributing to the prevention of CKD is the early identification of the presence of subclinical structural of functional sequelae after AKI episodes, due mainly to maladaptive repair mechanisms. These sequelae constitute a clear risk for the development of CKD, but many of them cannot be detected with traditional criteria based on the determination of pCr.. Hence the relevance of implementing new diagnostic methods based on biomarkers (identified in preclinical in vivo models) that allow the early monitoring of the pathophysiological mechanisms inherent to this maladaptive repair [25] [26].

The nephrotoxic and ischemic experimental models used in our study generate functional alterations characterized by a decrease in renal filtration (increases in pCr and blood urea nitrogen together with a decrease in CrCl, although within the ranges considered non-pathological). These results suggest an almost complete restoration of glomerular filtration capacity in both models. However, functional normalization does not imply complete renal recovery, as the histological analysis revealed persistent structural alterations, particularly fibrosis, even after apparent functional normalization. This dissociation between renal filtration and structure is consistent with previous studies highlighting the limitations of traditional biomarkers in detecting ongoing subclinical kidney damage [27]. In the ischemia-reperfusion model, tubular dilation, epithelial disruption, and the presence of cellular debris were evident on day 13, with partial resolution observed over the following week. However, ECM deposition increased during this period and persisted, particularly in the cortical region, one-month post-injury. Similarly, cisplatin-induced AKI resulted in epithelial disorganization, inflammation, and ECM deposition, with only partial recovery over time. Notably, the extent of structural recovery was markedly lower in the cisplatin group compared to the ischemia group. These observations suggest that while both types of injury result in persistent fibrotic remodeling, the degree and duration of such alterations may depend on the etiology of the initial insult. These results show that under these experimental conditions, both ischemic and nephrotoxic damage evolved to AKD, in which histological damage and apparently normal GFR (although with slight modifications in plasma urea, pCr and CrCl levels) occur in parallel.

Proteomic analysis of urine samples provided further insights into the molecular footprint of subclinical AKD. The distinct urinary proteomic profiles observed between cisplatin and I/R groups underscore the notion that the pathophysiological mechanisms underlying recovery and progression to fibrosis are injury-specific. Cisplatin-treated animals exhibited a more extensive proteomic alteration, with 44 differentially excreted proteins compared to controls, 33 of which were unique to this comparison. In contrast, only 15 proteins were differentially excreted in the I/R group relative to controls, with 6 being unique. Proteins exclusive to the cisplatin group were enriched in pathways related to coagulation, inflammation, and immune regulation—hallmarks of AKI-to-CKD transition and chronic kidney injury progression [28] [29]. Interestingly, we also identified 29 proteins commonly deregulated in both injury models (not found in controls), further supporting the concept of a shared subclinical AKD signature. These proteins were primarily associated with immune response and haemostasis, reinforcing their potential role in the maladaptive repair process following AKI.

Together, these findings suggest that GFR recovery following AKI may be misleading if considered in isolation, as subclinical structural and molecular alterations may persist and contribute to long-term adverse outcomes such as CKD. The proteomic signatures identified here may serve for the non-invasive and early detection of subclinical AKD and for distinguishing between injury etiologies. Moreover, the persistence of fibrosis despite apparent recovery of glomerular function highlights the need for therapeutic strategies aimed at mitigating fibrotic remodeling in the aftermath of AKI. Furthermore, our analysis has led to the identification of numerous urinary proteins involved in the previously described processes, many of which had not been previously associated with AKD. These findings serve as a foundation for future mechanistic studies aimed at elucidating the specific pathophysiological information conveyed by these proteins. They also pave the way for validation studies in patient populations to explore their potential utility as diagnostic and prognostic markers. Additionally, understanding the true role of these urinary proteins during the transition to CKD could open new avenues for the development of pharmacological strategies aimed at preventing or slowing CKD progression in its early stages. It is important to emphasize that the detection of subclinical kidney damage through the assessment of urinary biomarkers largely depends on the etiology of the initial AKI episode that has led to the development of AKD. Therefore, particular caution must be exercised when selecting appropriate diagnostic urinary biomarkers to identify the potential presence of subclinical renal injury. This selection should be tailored to the specific clinical context of each patient,particularly taking into account the underlying cause of the AKI episode, which may warrant close monitoring and further evaluation for possible hidden renal damage.

Interestingly, our findings also revealed a substantial number of urinary proteins that are consistently excreted regardless of the specific triggering factor behind AKI. This observation suggests the existence of a core proteomic response associated with AKD that is independent of the initial etiology. The identification of such non-specific yet consistently expressed proteins may hold significant diagnostic value, as these proteins could serve as more general biomarkers of AKD, enabling early detection even in cases where the primary insult is unclear or multifactorial. From a clinical perspective, this is especially relevant, as many patients present with overlapping or ambiguous clinical features that complicate the identification of the underlying cause, or have an unknown history of silent and undocumented AKIs. By focusing on these etiology-independent markers, it may be possible to develop more robust diagnostic tools that transcend the limitations of current etiologically-driven models. Furthermore, the presence of a common urinary proteomic signature in AKD raises important questions about shared downstream pathways of kidney damage and repair, potentially pointing to fundamental mechanisms of disease progression. Investigating the biological roles and regulatory networks of these proteins could yield new insights into the pathogenesis of AKD and its transition to CKD, ultimately contributing to the development of broader, more effective diagnostic and therapeutic strategies.

Understanding the principal biological processes associated with urinary proteomic fingerprints offers a valuable opportunity to deepen our knowledge of the pathophysiological mechanisms underlying AKD. The proteins identified through urinary proteomics can serve not only as diagnostic biomarkers but also as indicators of specific molecular pathways involved in renal injury, repair, and maladaptive responses [30] [31]. However, it is important to interpret these findings with caution. Although some proteins detected in urine may originate from renal tissues, it is plausible that others derive from the circulation. Changes in glomerular permeability or tubular handling may lead to the appearance of plasma proteins in the urine, particularly under pathological conditions where tubular reabsorption is impaired [31] [32]. For example, proximal tubular dysfunction can reduce the reabsorption of filtered proteins, leading to their increased urinary excretion, independent of their synthesis or release by renal cells [33]. Moreover, the urinary proteome is influenced by a range of physiological and pathological variables, including protein filtration, degradation, and reabsorption dynamics [34]. Therefore, the presence of specific proteins in urine may not always directly reflect local renal production, but rather altered renal handling due to structural or functional damage. Thus, while urinary proteomics provides critical insights into AKD, further validation—such as correlating urinary markers with tissue expression and functional studies—is essential to accurately delineate their origin and interpret their clinical relevance.

In summary, we have identified distinct urinary proteomic signatures linked to nephrotoxic and ischemic damage, as well as to the pathophysiological processes associated with progressive fibrosis of various etiologies. These findings form the basis of a novel diagnostic system that, in the near future, could facilitate early-stage monitoring of CKD and support the development of both preventive and palliative strategies.

## Acknowledgements

Joana Mercado-Hernández is recipient of a predoctoral fellowship from the Junta de Castilla y Leon (Spain) and the European Social Fund from the European Commission. The proteomic analysis was performed in the Proteomics Unit (CAI Biological Techniques) of Complutense University of Madrid.

## Author contributions

Joana Mercado-Hernández, Isabel Fuentes-Calvo, David Martín-Calvo, Sandra M. Sancho-Martínez: acquisition and analysis of data; Sandra M. Sancho-Martínez, Francisco J. López-Hernández and Carlos Martínez-Salgado: conception, design, interpretation of data, drafting of the work, and final approval of the version to be published.

## Data availability statement

The data underlying this article will be shared on reasonable request to the corresponding author.

## REFERENCES

1 Thomas ME, Blaine C, Dawnay A, Devonald MAJ, Ftouh S, Laing C, et al. The definition of acute kidney injury and its use in practice. Kidney Int. 2015 Jan;87(1):62–73.

2 Ronco C, Bellomo R, Kellum JA. Acute kidney injury. Lancet (London, England). 2019 Nov;394(10212):1949–64.

3 Siew ED, Davenport A. The growth of acute kidney injury: a rising tide or just closer attention to detail? Kidney Int. 2015 Jan;87(1):46–61.

4 Mehta RL, Cerdá J, Burdmann EA, Tonelli M, García-García G, Jha V, et al. International Society of Nephrology’s 0by25 initiative for acute kidney injury (zero preventable deaths by 2025): a human rights case for nephrology. Lancet (London, England). 2015 Jun;385(9987):2616–43.

5 Lewington AJP, Cerdá J, Mehta RL. Raising awareness of acute kidney injury: a global perspective of a silent killer. Kidney Int. 2013;84(3):457–67.

6 Melo FDAF, Macedo E, Bezerra ACF, De Melo WAL, Mehta RL, Burdmann EDA, et al. A systematic review and meta-analysis of acute kidney injury in the intensive care units of developed and developing countries. PLoS One. 2020 Jan;15(1). DOI: 10.1371/JOURNAL.PONE.0226325

7 Kellum JA, Sileanu FE, Bihorac A, Hoste EAJ, Chawla LS. Recovery after Acute Kidney Injury. Am J Respir Crit Care Med. 2017 Mar;195(6):784–91.

8 Forni LG, Darmon M, Ostermann M, Oudemans-van Straaten HM, Pettilä V, Prowle JR, et al. Renal recovery after acute kidney injury. Intensive Care Med. 2017 Jun;43(6):855–66.

9 Zuk A, Bonventre J V. Acute Kidney Injury. Annu Rev Med. 2016 Jan;67:293–307.

10 Bihorac A, Yavas S, Subbiah S, Hobson CE, Schold JD, Gabrielli A, et al. Long-term risk of mortality and acute kidney injury during hospitalization after major surgery. Ann Surg. 2009 May;249(5):851–8.

11 Nejat M, Pickering JW, Devarajan P, Bonventre J V., Edelstein CL, Walker RJ, et al. Some biomarkers of acute kidney injury are increased in pre-renal acute injury. Kidney Int. 2012 Jun;81(12):1254–62.

12 Heung M, Chawla LS. Acute kidney injury: gateway to chronic kidney disease. Nephron Clin Pract. 2014 Apr;127(1–4):30–4.

13 Haase M, Kellum JA, Ronco C. Subclinical AKI--an emerging syndrome with important consequences. Nat Rev Nephrol. 2012 Dec;8(12):735–9.

14 de Geus HR, Haase M, Jacob L. The cardiac surgery-associated neutrophil gelatinase-associated lipocalin score for postoperative acute kidney injury: Does subclinical acute kidney injury matter? J Thorac Cardiovasc Surg. 2017 Sep;154(3):939–40.

15 Chawla LS, Eggers PW, Star RA, Kimmel PL. Acute kidney injury and chronic kidney disease as interconnected syndromes. N Engl J Med. 2014 Jul;371(1):58–66.

16 Lameire NH, Levin A, Kellum JA, Cheung M, Jadoul M, Winkelmayer WC, et al. Harmonizing acute and chronic kidney disease definition and classification: report of a Kidney Disease: Improving Global Outcomes (KDIGO) Consensus Conference. Kidney Int. 2021 Sep;100(3):516–26.

17 Taylor KM, Au AYM, Herath S, Succar L, Wong J, Erlich JH, et al. Kidney functional reserve and damage biomarkers in subclinical chronic kidney disease and acute kidney injury. Am J Physiol Renal Physiol. 2023 Dec;325(6):F888–98.

18 Lopez-Novoa JM, Quiros Y, Vicente L, Morales AI, Lopez-Hernandez FJ. New insights into the mechanism of aminoglycoside nephrotoxicity: an integrative point of view. Kidney Int. 2011;79(1):33–45.

19 Cuesta C, Fuentes-Calvo I, Sancho-Martinez SM, Valentijn FA, Düwel A, Hidalgo-Thomas OA, et al. Urinary KIM-1 Correlates with the Subclinical Sequelae of Tubular Damage Persisting after the Apparent Functional Recovery from Intrinsic Acute Kidney Injury. Biomedicines. 2022 May;10(5). DOI: 10.3390/BIOMEDICINES10051106

20 Fuentes-Calvo I, Cuesta C, Sancho-Martínez SM, Hidalgo-Thomas OA, Paniagua-Sancho M, López-Hernández FJ, et al. Biomarkers of persistent renal vulnerability after acute kidney injury recovery. Sci Rep. 2021 Dec;11(1). DOI: 10.1038/S41598-021-00710-Y

21 Casanova AG, Vicente-Vicente L, Hernández-Sánchez MT, Prieto M, Rihuete MI, Ramis LM, et al. Urinary transferrin pre-emptively identifies the risk of renal damage posed by subclinical tubular alterations. Biomed Pharmacother. 2020 Jan;121. DOI: 10.1016/J.BIOPHA.2019.109684

22 Yin C, Wang N. Kidney injury molecule-1 in kidney disease. Ren Fail. 2016 Nov;38(10):1567–73.

23 Pellicer-Valero ÓJ, Massaro GA, Casanova AG, Paniagua-Sancho M, Fuentes-Calvo I, Harvat M, et al. Neural Network-Based Calculator for Rat Glomerular Filtration Rate. Biomedicines. 2022 Mar;10(3). DOI: 10.3390/BIOMEDICINES10030610

24 Szklarczyk D, Gable AL, Lyon D, Junge A, Wyder S, Huerta-Cepas J, et al. STRING v11: protein-protein association networks with increased coverage, supporting functional discovery in genome-wide experimental datasets. Nucleic Acids Res. 2019 Jan;47(D1):D607–13.

25 Fiorentino M, Grandaliano G, Gesualdo L, Castellano G. Acute Kidney Injury to Chronic Kidney Disease Transition. Contrib Nephrol. 2018;193:45–54.

26 Charlton JR, Li T, Wu T, deRonde K, Xu Y, Baldelomar EJ, et al. Use of novel structural features to identify urinary biomarkers during acute kidney injury that predict progression to chronic kidney disease. BMC Nephrol. 2023 Dec;24(1). DOI: 10.1186/S12882-023-03196-0

27 Bonventre J V., Vaidya VS, Schmouder R, Feig P, Dieterle F. Next-generation biomarkers for detecting kidney toxicity. Nat Biotechnol. 2010 May;28(5):436–40.

28 Guzzi F, Cirillo L, Roperto RM, Romagnani P, Lazzeri E. Molecular Mechanisms of the Acute Kidney Injury to Chronic Kidney Disease Transition: An Updated View. Int J Mol Sci. 2019 Oct;20(19). DOI: 10.3390/IJMS20194941

29 Franzin R, Stasi A, Fiorentino M, Stallone G, Cantaluppi V, Gesualdo L, et al. Inflammaging and Complement System: A Link Between Acute Kidney Injury and Chronic Graft Damage. Front Immunol. 2020 May;11. DOI: 10.3389/FIMMU.2020.00734

30 Mishra J, Ma Q, Kelly C, Mitsnefes M, Mori K, Barasch J, et al. Kidney NGAL is a novel early marker of acute injury following transplantation. Pediatr Nephrol. 2006 Jun;21(6):856–63.

31 Vaidya VS, Waikar SS, Ferguson MA, Collings FB, Sunderland K, Gioules C, et al. Urinary biomarkers for sensitive and specific detection of acute kidney injury in humans. Clin Transl Sci. 2008 Dec;1(3):200–8.

32 Schaub S, Wilkins JA, Antonovici M, Krokhin O, Weiler T, Rush D, et al. Proteomic-based identification of cleaved urinary beta2-microglobulin as a potential marker for acute tubular injury in renal allografts. Am J Transplant. 2005 Apr;5(4 Pt 1):729–38.

33 Sancho-Martínez SM, Blanco-Gozalo V, Quiros Y, Prieto-García L, Montero-Gómez MJ, Docherty NG, et al. Impaired Tubular Reabsorption Is the Main Mechanism Explaining Increases in Urinary NGAL Excretion Following Acute Kidney Injury in Rats. Toxicol Sci. 2020 May;175(1):75–86.

34 Zhou Y, Vaidya VS, Brown RP, Zhang J, Rosenzweig BA, Thompson KL, et al. Comparison of kidney injury molecule-1 and other nephrotoxicity biomarkers in urine and kidney following acute exposure to gentamicin, mercury, and chromium. Toxicol Sci. 2008 Jan;101(1):159–70.

