## Supplemental Figures for "SUBCLINICAL ACUTE KIDNEY DISEASE: URINARY PROTEOMIC DIVERGENCE FOLLOWING NEPHROTOXIC AND ISCHAEMIC INJURY"

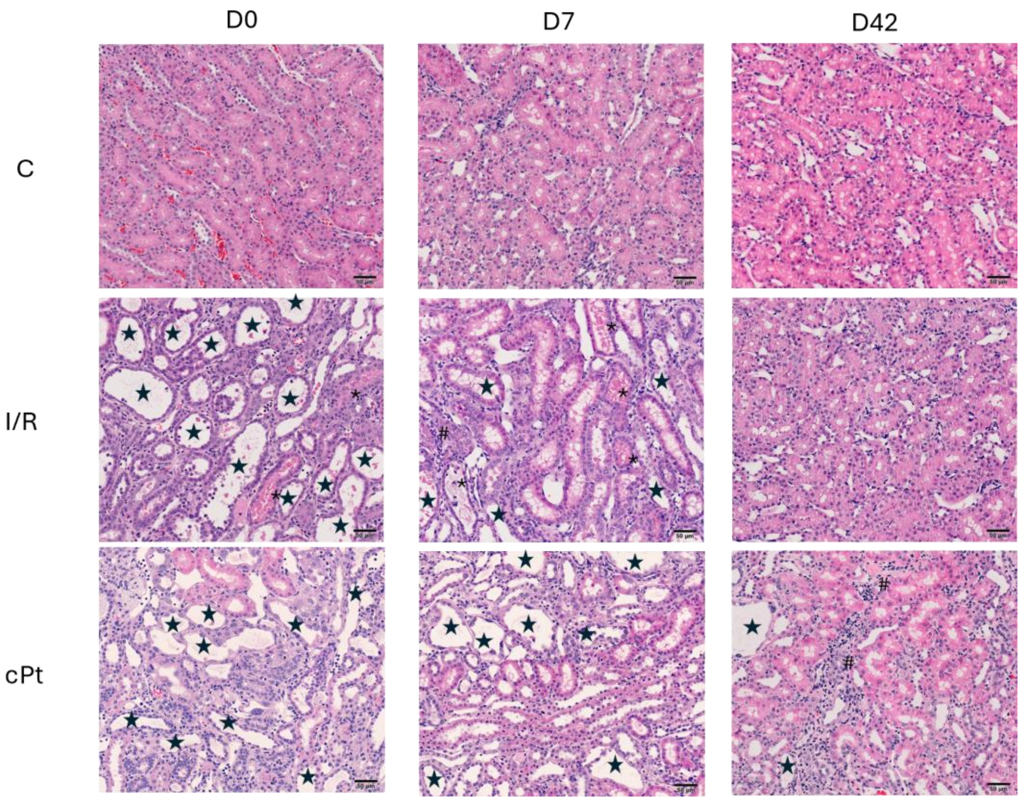


**Supplementary Figure 1**. **Renal histology.** Representative images of renal outer medulla specimens stained with haematoxylin and eosin (HE) at days 0, 7 and 42 (magnification: 200X). Casts and cellular debris (*), tubular atrophy (star) and inflammatory infiltrates (#) are visible in both AKI groups, both in the cortex and in the outer medulla. This damage gradually decreases until day 42, where only isolated lesions are observed in the cPt group. C: control; cPt: cisplatin group; I/R: ischaemia-reperfusion group; D0, D7, D42: days 0, 7 and 42 respectively.

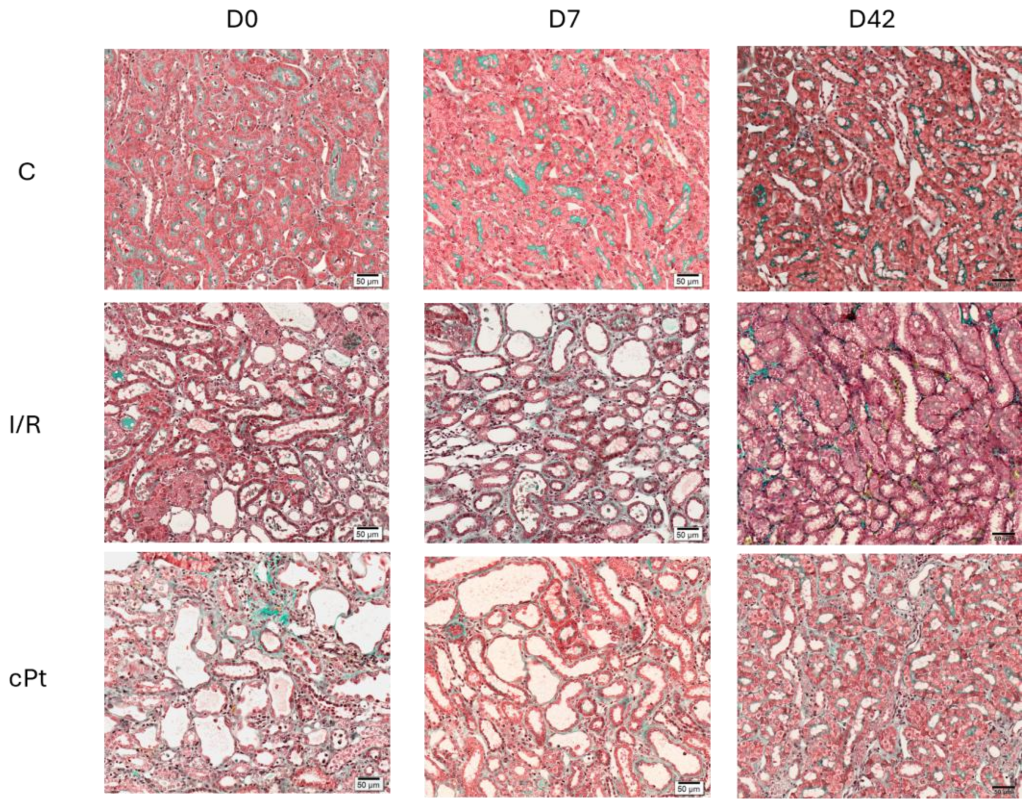


**Supplementary Figure 2**. **Fibrosis assessment**. Representative images of renal outer medulla  specimens stained with Masson trichrome at days 0, 7 and 42; magnification: 200X. There is a progressive fibrosis marked by the accumulation of collagen fibers (stained in green). C: control; cPt: cisplatin group; I/R: ischaemia-reperfusion group; D0, D7, D42: days 0, 7 and 42 respectively.
