## Supplementary material for "SUBCLINICAL ACUTE KIDNEY DISEASE: URINARY PROTEOMIC DIVERGENCE FOLLOWING NEPHROTOXIC AND ISCHAEMIC INJURY": Tables

Table 1. Proteins excreted in the cisplatin group, but absent in control and ischaemia groups.

| Uniprot Accesion Number | Protein description |
|---|---|
| A0A0G2JSI5 | Chymotrypsin-like elastase family member 1 |
| B2RYM3 | Inter-alpha trypsin inhibitor, heavy chain 1 |
| D4A499 | Phospholipase A2 inhibitor and LY6/PLAUR domain-containing |
| P04636 | Malate dehydrogenase, mitochondrial |
| P14141 | Carbonic anhydrase 3 |
| Q5BJT9 | Creatine kinase |
| Q5I0D7 | Xaa-Pro dipeptidase |
| Q7TQ70 | Fibrinogen alpha chain |
| Q9QZQ5 | CCN family member 3 |

Table 2. Proteins excreted in the ischaemia group, but absent in control and cisplatin groups.

| Uniprot Accesion Number | Protein description |
|---|---|
| D4ABU7 | Triggering receptor expressed on myeloid cells 1 |
| F1LYU4 | Ig-like domain-containing protein |
| F1M698 | CD209f antigen |
| M0RC23 | Ig-like domain-containing protein |
| P46720 | Solute carrier organic anion transporter family member 1A1 |
| P63095 | Guanine nucleotide-binding protein G(s) subunit alpha isoforms short |
| Q6AYT0 | Quinone oxidoreductase |
| Q80W57 | Broad substrate specificity ATP-binding cassette transporter ABCG2 |

Table 3. Proteins excreted in cisplatin + ischaemia groups, but absent in the control group

| Uniprot Accesion Number | Protein description | Fold Change (log2): (Cpt) / (Isq) | FDR: (Cpt) / (Isq) |
|---|---|---|---|
| A0A0G2K458 | Ig-like domain-containing protein | -0.897802556 | 0.695763069 |
| A0A0G2K6T9 | Protocadherin 1 | -0.036654919 | 0.974990712 |
| A0A0G2K7I1 | Ig-like domain-containing protein | 0.111830077 | 0.952361276 |
| B1H259 | Cochlin | 1.503164109 | 0.198605785 |
| D3ZBB2 | Ig-like domain-containing protein | -0.213172966 | 0.899736527 |
| D3ZDK4 | Angiopoietin-like 7 | 2.184077583 | 0.079687458 |
| D3ZEP5 | Ig-like domain-containing protein | 0.346471139 | 0.819888598 |
| D3ZY96 | Neutrophilic granule protein | -0.586701374 | 0.691426399 |
| D4A0W2 | Lysozyme f1 | 0.799823118 | 0.557494531 |
| D4A6I7 | Prostate stem cell antigen | -1.374256488 | 0.364610406 |
| G3V7P2 | Fibrinogen-like 2 | 0.70549757 | 0.459427646 |
| G3V8V1 | Granulin, isoform CRA_c | -1.470147993 | 0.488288219 |
| M0R9U2 | Ig-like domain-containing protein | 0.261398525 | 0.860439231 |
| P02631 | Oncomodulin | 1.187803925 | 0.117034948 |
| P02680 | Fibrinogen gamma chain | 1.766304839 | 0.17676259 |
| P04639 | Apolipoprotein A-I | 1.751888656 | 0.203963307 |
| P08753 | Guanine nucleotide-binding protein G(i) subunit alpha | -1.353245661 | 0.488288219 |
| P11517 | Hemoglobin subunit beta-2 | 2.065479319 | 0.463427741 |
| P11883 | Aldehyde dehydrogenase, dimeric NADP-preferring | -0.070540789 | 0.963400415 |
| P14480 | Fibrinogen beta chain | 2.120144024 | 0.19337314 |
| P17559 | Uteroglobin | 1.585201518 | 0.29766106 |
| P36375 | Glandular kallikrein-10 | -1.17144615 | 0.557533664 |
| P80202 | Activin receptor type-1B | -0.984498309 | 0.664495829 |
| Q06496 | Sodium-dependent phosphate transport protein 2A | -1.346139876 | 0.246428843 |

| | | | |
|---|---|---|---|
| Q4QQV8 | Charged multivesicular body protein 5 | -2.225199096 | 0.052685042 |
| Q6AYE5 | Out at first protein homolog | 0.385947514 | 0.752130459 |
| Q6VPP3 | Chloride channel accessory 4 | 2.741343858 | 0.37245626 |
| Q9EQV9 | Carboxypeptidase B2 | 1.884228317 | 0.172630871 |
| Q9R1T3 | Cathepsin Z | 1.565797004 | 0.052685042 |

Table 4. Proteins excreted in the control group, but absent in cisplatin and ischaemia groups.

| Uniprot Accesion Number | Protein description |
|---|---|
| B2RYC9 | Glucosylceramidase |
| G3V862 | Angiopoietin-like 2 |
| M0R7T8 | Similar to Vomeromodulin |
| Q80WD1 | Reticulon-4 receptor-like 2 |
| Q9Z2L0 | Voltage-dependent anion-selective channel protein 1 |
